# Essential oils of plants and their combinations as an alternative adulticides against *Anopheles gambiae* (Diptera: Culicidae) populations

**DOI:** 10.1101/2022.05.01.490040

**Authors:** Mahamoudou Balboné, Ignace Sawadogo, Dieudonné Diloma Soma, Samuel Fogné Drabo, Moussa Namountougou, Koama Bayili, Rahim Romba, Georges Benson Meda, Honorat Charles Roger Nebié, Roch K. Dabire, Imae□l H. N. Bassolé, Olivier Gnankine

**Affiliations:** Laboratoire d’Entomologie Fondamentale et Appliquée, Unité de Formation et de Recherche en Sciences de la Vie et de la Terre (UFR-SVT), Université Joseph KI-ZERBO, Ouagadougou, 03 BP 7021, Burkina Faso; Institut de Recherche en Sciences Appliquées et Technologies, Ouagadougou 03 BP 7047, Burkina Faso; Institut de Recherche en Sciences de la Santé/Centre Muraz, Bobo-Dioulasso BP 545, Burkina Faso; Université Nazi Boni, Bobo-Dioulasso, Burkina Faso

**Keywords:** Resistance, essential oil, combination, major compound, mortalities, *Anopheles gambiae*

## Abstract

The persistence of malaria and the increasing of resistance of *Anopheles gambiae* species to chemicals remain major public health concerns in sub-Saharan Africa. Faced to these concerns, the search for alternative vector control strategies as use of essential oils need to be implemented. Here, the five essential oils (EOs) from *Cymbopogon citratus, Cymbopogon nardus, Eucalyptus camaldulensis, Lippia multiflora, Ocimum americanum* obtained by hydro distillation were tested according to WHO procedures on *Anopheles gambiae* “Kisumu” and field strains collected in “Vallée du Kou”. Also, the binary combinations of *C. nardus* and *O. americanum* were examined. As results, among the EOs tested, *L. multiflora* was the most efficient regarding KDT_50_ and KDT_95_ and rate of morality values. Our current study showed that C8 (C.n 80% : O.a 20%) and C9 (C.n 90% : O.a 10%), were more toxic to *Anopheles gambiae* strain VK with the mortality rates reaching 80.7 and 100% at the 1% concentration, respectively. These two binary combinations shown a synergistic effect on the susceptible population. However, only C9 showed a synergistic effect on VK population. The bioactivity of the two EOs, *Cymbopogon nardus* and *Ocimum americanum*, was improved by the combinations at certain proportions and may constitute an alternative to pyrethroid resistance.

## BACKGROUND

Malaria remains one of the world’s most deadliest diseases. It is a life-threatening disease caused by parasites that are transmitted to people through the bites of infected female Anopheles mosquitoes. Each year, 154 to 289 million persons are infected with 490–836 thousand deaths recorded mostly in children under 5. About 90% of this burden is recorded in Africa^1^. In Burkina Faso, malaria is a major health issue and accounts for 43% of medical consultations and 22% of deaths were recorded. This country is among the 10 most affected (3% of cases and 4% of deaths worldwide)^2^ by malaria disease.

Until now, long-lasting insecticidal nets (LLINs) and indoor residual spraying (IRS) remain the two main tools commonly used in controlling adults of *Anopheles gambiae*, the vector of malaria^3^. Pyrethroids are only products used for LLINs^4^. However, pyrethroid resistance is a threat to the efficiency of these protective tools, especially when resistance is occurred at high levels^5,6^. In Burkina Faso, pyrethroid resistance has been observed throughout the country for several years and there is concern that pyrethroid-based LLINs does not provide the expected levels of individual and community protection^7^.

To improve the effectiveness of vector control tools, the World Health Organization (WHO) has developed a global plan for the management of insecticide resistance (GPIRM)^8^. Key elements of this plan include: (i) insecticide rotation; (ii) mixtures of at least two different insecticides; (iii) alternate use of at least two insecticides of different classes; and, (iv) mosaic use of insecticides. Today, a limiting factor in the development of these strategies were the absence of alternative classes of insecticides for LLINs. However, in recent years, several studies have shown that the use of piperonyl butoxide (PBO) as a synergist restores susceptibility to deltamethrin or permethrin in several regions of the country^9-11^.

To manage the resistance of insects generally and mosquitoes specifically to chemical insecticides, many research programmes have focused on natural products derived from plants as an alternative to conventional insecticides used in vector control for which resistance has been detected^12^. Among the many natural products, essential oils (EOs) and their constituents have received considerable attention in the search for new pesticides and have been found to possess insecticidal properties^13^.

Essential oils (EOs) from plants are secondary metabolites comprising different bioactive compounds and have gained importance in terms of alternative to chemicals. They are biodegradable, environmentally safe and easy to use and compose of a mixture of different bioactive compounds which offer less chance for emerging resistance^14^.

Previous studies have identified more than 3,000 compounds from 17,500 aromatic plants ^15^. Most of them have been tested for their insecticidal properties and have been reported to have insecticidal effects^16-17^. Other studies have shown equal or higher toxicity of the major compounds compared to its whole EO^18^. Some studies have also attempted to formulate efficient insecticides by combining different plants with chemical insecticides to increase the overall toxicity and minimize the secondary effects^19^. Recent studies have shown insecticidal properties of EO of *Cymbopogon nardus, Eucalyptus camaldulensis, Lippia multiflora*, and *Ocimum americanum*^20^. These studies focused on *Aedes aegypti* populations in the city of Ouagadougou, Central Burkina Faso.

The objective of the current study aimed at examining the adulticidal activities of essential oils of five aromatic plants of *Cymbopogon citratus, Cymbopogon nardus, Eucalyptus camaldulensis, Lippia multiflora and Ocimum americanum*, on *Anopheles* populations collected in “Vallée du Kou” (Bama), Western Burkina Faso. This current study, also evaluate the toxicity level of binary combinations of essential oils of *C. nardus* and *O. americanum*, on *An. gambiae* populations.

## Materials and Methods

### Sampling of larvae and rearing

Larvae and pupae of the resistant strain of *Anopheles gambiae* “VK” were collected from June to October 2021, in the “Vallée du Kou” (Bama). The “Vallée du Kou” (VK) is a district of the Bama department (11° 23’ 59’’ N, 4° 25’ 46’’ W) in the province of Houet, the economic capital of Burkina Faso. The larvae were brought and reared in the insectarium of the “Institut de Recherches en Sciences de la Santé/Direction Régionale de l’Ouest” (IRSS/DRO) located in Bobo-Dioulasso. The larvae were fed with tetraMin (Tetrawerke, Melle, Germany). Adult mosquitoes emerging from the pupae of the collected strain were placed in cages and fed with 10% sugar solution. Female mosquitoes of the resistant strain were used for susceptibility tests. It is the same for the susceptible strain of *An. gambiae* “Kisumu” maintained at the insectarium, and used as a reference strain in this current study.

### Essential oils extraction

The essential oils were obtained from the five aromatic plant species, *Cymbopogon citratus, Cymbopogon nardus* (Linn.), *Eucalyptus camaldulensis, Lippia multiflora*, and *Ocimum americanum*. They were extracted at the “Institut de Recherche en Sciences Appliquées et Technologies” (IRSAT) by hydrodistillation (HD) using a *clevenger*-type apparatus and stored in a dark glass bottle at 4 °C prior to use. The combinations of two EOs were made, after having done the bioassay tests with the whole EOs.

### Analysis by gas chromatography/flame ionization detector (GC/FID)

Gas chromatography/flame ionization detector (GC-FID) analysis of essential oils of *C. citratus, C. nardus, E. camaldulensis, L. multiflor*a and *O. americanum* obtained from their leaves was performed on an Agilent 6890N GC instrument equipped with a FID, with a narrow bore DB-5 column (length 10 m, inner diameter 0.1 mm, film thickness 0.17 mm; Agilent, Palo Alto, CA). The oven temperature was programmed from 60 °C to 165 °C at 8 °C/min and from 165 °C to 280 °C at 20 °C/min, with 1 minute of post-operation at 280 °C. Diluted samples (1/100 in sample) were subjected to an ionization test. Diluted samples (1/100 in acetone) of 1.0 µl were injected manually and without fractionation. The percentage peak area was calculated on the basis of the FID signal using the GC HP-Chemstation software (Agilent Technologies).

### Gas chromatography/mass spectrometry (GC/MS) analysis

GC/MS analysis was performed on a GC HP 6890 coupled to a MSD HP 5972 (Hewlett Packard, Palo Alto, CA), and was equipped with a ZB-5MS Zebron capillary column (length 30 m, ID 0.25 mm, film thickness 0.25 mm; Agilent). The carrier gas used was helium and the oven temperature were maintained at 45 °C for 2 min and then increased from 45 °C to 165 °C (4 °C/min) and then from 165 °C to 280 °C (15 °C/min).

### Bioassays on adult mosquitoes

Susceptibility tests were carried out using WHO insecticide susceptibility test-kits and standard procedures^3^. Impregnation of Whatman n°1 papers were done according to the protocol adopted by N’Guessan et al^21^. Four rectangular papers (size 12 cm× 15 cm) were impregnated with 2 ml of a given concentration of an essential oil/combination of EOs diluted in acetone at varying proportions. Three concentrations, 0.1%, 0.5% and 1%, were used for each EO or combination of *Cymbopogon nardus* (C.n) and *Ocimum americanum* (O.a).

For combination of EOs and for each concentration (0.1; 0.5 and 1%), 9 binary combinations: i) C.n 10% : O.a 90%; ii) C.n 20% : O.a 80%; iii) C.n 30% : O.a 70%; iv) C.n 40% : O.a 60 %; v) C.n 50% : O.a 50%; vi) C.n 60% : O.a 40%; vii) C.n 70% : O.a 30%; viii) C.n 80% : O.a 20% and ix) C.n 90% : O.a 10% corresponding to C1, C2, C3, C4, C5, C6, C7, C8 and C9, respectively were done. Control papers were impregnated with 2 ml of acetone only. Tests were carried out at 25°C (± 2°C) and 70-80% relative humidity. The number of mosquitoes knocked down was recorded every 5 minutes. Permethrin 0.75% was used as positive control insecticide. After the exposure time, mosquitoes were transferred to holding tubes and were fed with 10% sugar juice for 24 hours. Subsequently the mortality was recorded. The susceptible strain *Anopheles gambiae* “Kisumu” was used as reference to determine the diagnostic doses.

### Data analysis

The data obtained from the bioassays performed were analyzed using XLSTAT statistical software version 2015.1.01. The knock-down time (KDT_50_ and KDT_95_), lethal concentrations (LC_50_ and LC_99_) and 95% confidence limits (95% CL) were calculated by probity analysis using the same statistical software in order to compare the toxicity of the plant essential oils against the tested mosquito adults. The KDT_50_, KDT_95_, CL_50_, CL_99_ values and mortalities were considered significantly different between the essential oils (p < 0.05) if the 95% CL (Confident Limits) did not overlap. In all tests, no control mortality was detected after the 24-hour exposure; therefore, no correction was required based on Abbot’s formula^22^.

Interactions between the combinations performed were determined using the Fractional Inhibitory Concentration indices or FIC indices. These indices were calculated in the following way: FIC indice = FIC_A_+ FIC_B_ and FIC_A_ = MIC_AB_ / MIC_A_ and FIC_B_ = MIC_AB_ / MIC_B_ where FIC_A_ and FIC_B_ are the minimum inhibitory concentrations that kill adult mosquitoes for EO A and B respectively.

Thus, we have i) FICA: Fractional Concentration of A; ii) FICB: Fractional Concentration of B; iii) MICAB: Minimum Inhibitory Concentration of A in the combination; iv) MICA: Minimum Inhibitory Concentration of A; v) MICAB: Minimum Inhibitory Concentration of B in the combination.

The results were interpreted as follows: i) Synergy: FIC < 0.5; ii) additive: 0.5 ≤ FIC ≤ 1; iii) indifferent: 1 ≤ FIC ≤ 4; or iv) antagonism: FIC > 4^23,24^.

## RESULTS

### Chemical composition of the essential oils

The main compounds of the 5 essential oils are summarized in Table 1. The EO of *C. citratus*, consisting only of oxygenated monoterpenes (99.9%) which were neral (44.7%) and geranial (55.2%). The EO of *C. nardus* consisted of six compounds mostly oxygenated monoterpenes (77.9%), characterized by citronellal (41.7%), geraniol (20.8%) and β-elemene (11%). As for EO of *E. camaldulensis*, it consisted of sixteen compounds mainly hydrocarbon monoterpenes, rich in 1,8-cineole (59.5%). The EO of *L. multiflora* consisted of sixteen compounds dominated by monoterpenes (69.51%), characterized by β-Caryophyllene (20.1%), p-cymene (14.6%), thymol acetate (12.0 %) and 1.8 cineol (11.6%) whereas that of *O. americanum* consisted of twenty-six (26) compounds characterized by a high percentage of 1.8-cineole (31.22%) followed by camphor (12.73%).

**TABLE 1:** Major compounds of the 5 essential oils tested on adults of *Anopheles gambiae*

| Essential oils | Major compounds | Retention indices | Mono Hydro (%) | Mono Oxy (%) | Sesqui Hydro (%) | Sesqui Oxy (%) |
| --- | --- | --- | --- | --- | --- | --- |
| *C. citratus* | Geranial | 1268 | – | 55.2 | – | – |
|  | Neral | 1242 | – | 44.7 | – | – |
|  | **Total other compounds** |  | - | - | - | – |
| *C. nardus* | Citronellal | 1158 | – | 41.7 | – | – |
|  | Geraniol | 1253 | – | 20.8 | – | – |
| | β-Elemene | 1372 | – | – | 11 | – |
|  | **Total other compounds** |  | – | 15.4 | 3.7 | – |
| *E. camaldulensis* | 1,8 cineol | 1034 | – | 59.55 | – | – |
|  | **Total other compounds** |  | 24.89 | 7.41 | 1.4 | 0.34 |
| *L. multiflora* | β-caryophyllene | 1415 | – | – | 20.1 | - |
|  | p-cymene | 1027 | 14.6 | – | – | - |
|  | Thymul acetate | 1355 | – | 12 | – | - |
|  | 1.8 cineol | 1034 | - | 11.6 | - | - |
|  | **Total other compounds** |  | 18.41 | 12.9 | 5.7 | 4.5 |
| *O. americanum* | 1,8 cineol | 1034 | – | 31.22 | – | – |
|  | Camphor | 1151 | – | 12.73 | – | – |
|  | **Total other compounds** |  | 29.59 | 43.95 | 13.03 | 0.99 |

Mono = monoterpenes, Sesqui = sesquiterpenes, Hydro = Hydrocarbon and oxy = Oxygenated

### Knock Down Times (KDT_50_ and KDT_95_) values

The KDT_50_ and KDT_95_ calculated with 95% confidence limits (CL) obtained from the essential oils tested and EO combinations of *C. nardus* and *O. americanum* are shown in tables 1 and 2, respectively. All EOs and combinations exhibited KD effects on both strains.

**Table 2:** KDT_50_, KDT_95_ and rates of mortality of essential oils tested on the susceptibility (Kisumu) and field (VK) strains of *Anopheles gambiae*.

| Essential oils | Doses (%) | *Anopheles gambiae* « Kisumu » |  |  | *Anopheles gambiae* « VK » |  |  |
| --- | --- | --- | --- | --- | --- | --- | --- |
|  |  | KDT<sub>50</sub> (min) | KDT<sub>95</sub> (min) | Mortality (%) | KDT<sub>50</sub> (min) | KDT<sub>95</sub> (min) | Mortality (%) |
| *C. citratus* | 0.1 | 103.24 | 204.66 | 11.05 | 140.2 | 318.7 | 2.15 |
|  | 0.5 | 77.10 | 155.86 | 17.35 | 124.9 | 275.8 | 3.33 |
|  | 1 | 60.19 | 132.39 | 35.29 | 116.5 | 233.6 | 4.76 |
| *C. nardus* | 0.1 | 88.37 | 196.99 | 18.48 | 148.6 | 241.8 | 2.15 |
|  | 0.5 | 33.3 | 108.9 | 26.09 | 90.2 | 149.5 | 15.21 |
|  | 1 | 5.8 | 11.5 | 86.36 | 12.4 | 70.7 | 40.63 |
| *E. camaldulensis* | 0.1 | 192.2 | 289.7 | 1.05 | 227.3 | 309.7 | 0 |
|  | 0.5 | 157.1 | 275.1 | 6.45 | 189.9 | 286.7 | 1.12 |
|  | 1 | 144.8 | 266.5 | 9.78 | 166.1 | 280 | 3.16 |
| *L. multiflora* | 0.1 | 19.8 | 58.8 | 30.30 | 87.2 | 142.7 | 13.16 |
|  | 0.5 | 10.5 | 19 | 90 | 32 | 57.7 | 33.56 |
|  | 1 | -28.3 | -8.2 | 100 | -5.9 | 1.5 | 96.88 |
| *O. americanum* | 0.1 | 137.5 | 225.6 | 1.10 | 156.4 | 267.5 | 0 |
|  | 0.5 | 87.06 | 146.64 | 6.73 | 144.7 | 248.9 | 2.2 |
|  | 1 | 30.04 | 60.76 | 41.49 | 116.3 | 204.6 | 6.38 |
| Permethrin | 0,75 | 16.3 | 30 | 100 | 41.2 | 90.4 | 62.5 |

KDT = Knock-down times

Using the essential oils alone on “VK” strain, the lowest KDT_50_ and KDT_95_ values were obtained with *L. multiflora* (87.2 and 142.7 min at 0.1%, 32 and 57.7 min at 0.5% respectively) followed by *C. nardus* (90.2 and 149.5 min at 0.5% and 12.4 and 70.7 min at 1%, respectively), by *O. americanum* (116.3 and 204.6 min at 1%), and *E. camaldulensis* (166.1 and 280 min at 1%). The KDT_50_ and KDT_95_ values of *C. citratus* were 116.3 and 233.6 min at 1%. The KDT values obtained with *L. multiflora* were significantly lower than those of permethrin 0.75% (KDT_50_ = 41.2 min; KDT_95_ = 90.4 min for permethrin 0.75 % on VK strain). The KDT effects for all OEs on the VK strain were significantly higher than those found with Kisumu strain.

Using the combinations on the VK field strain, the lowest KDT50 and KDT95 values were obtained with C9 (C.n 90% : O.a 10%) (53.5 and 126.7 min at 0.1%, -5 and 7.4 min at 0.5%, -227.3 and -69.6 min at 1% respectively) followed by C5 (C. n 50% : O.a 50%) (106.7 and 182.8 min at 0.1%, 23.5 and 47.7 min at 0.5% and 7.1 and 16.5 min at 1%, respectively) and C8 (C.n 80% : O.a 20%) (119 and 218.2 min, at 0.1%, 24.2 and 52.7 min at 0.5%, -2.1 and 12.2 min at 1%, respectively). On the same strain, the highest KDT50 and KDT95 values were obtained with C1 (C.n 10% : O.a 90%) (171.6 and 321.3 min at 0.1%, 33 and 57.3 min at 0.5% and 27.4 and 50.3 min at 1%, respectively) and C2 (C.n 10% : O.a 90%) (222 and 310 min at 0.1%, 35 and 73.3 min at 0.5%, 16 and 25.9 min at 1%, respectively).

On the Kisumu strain, the KDTs were significantly higher than those obtained on the VK strain, except with the C4 (C.n 40% : O.a 60%), C6 (C.n 60% : O.a 40%), C7 (C.n 70% : O.a 30%), C8 and C9 combinations at the highest concentration 1%. Also, on both *An. gambiae* strains, the KDT values with the combinations were significantly lower than those obtained with the two EOs used individually and lower than those obtained with permethrin 0.75%.

### Rates of Mortality and lethal concentrations (LC)

All EOs exhibited adulticidal activity on both strains. The rates of mortality of field strain of *An. gambiae* “VK” varied from 0.0% to 13.16% at 0.1%, from 1.12% to 23.56% at 0.5% and from 3.16% to 96.9 at 1% (Table 2). The highest rates of mortality were obtained with *L. multiflora* (33.56 and 96.88% at 0.5 and 1% concentrations, respectively), followed by *C. nardus* (40.68% at 1% concentrations) and the lowest rates of mortality were found with *E. camaldulensis* (3.16% at 1% concentration). Rate of Mortality was 6.38% at 1% concentrations for *O. americanum*, while it reached 62.5% for permethrin 0.75%. On Kisumu strain, rates of mortality were 100% and 86.36% at 1% concentration with *L. multiflora* and *C. nardus* EOs, respectively. There was significant difference between rates of mortality obtained on the two strains from *An. gambiae* except for *L. multiflora* EO at the 1% concentration.

Using combinations against *An. gambiae*, rates of mortality on the field strain of *An. gambiae* collected in VK ranged from 1.15 to 12.15%, 5.21 to 94% and 9.62 to 100% at concentrations 0.1, 0.5 and 1%, respectively (Tables 3). The combinations as C8 (C.n 80%: O.a 20%) at 1% and C9 (C.n 90% : O.a 10%) at 0.5 and 1% were toxic to VK populations regarding the rate of mortality that were above 80%. Indeed, on Kisumu strain, the combination of C9 (C.n 90% : O.a 10%) exhibits the same rate of mortality (100%) as found with permethrin. However, on VK strain, the C9 combination at 0.5% concentration showed significantly higher mortalities than those of permethrin.

**Table 3:** KDT_50_, KDT_95_ and rates of mortality of essential oils combinations of *C. nardus* (C.n) and

| Codes | Combination of essential oil | Doses (%) | *Anopheles gambiae* "Kisumu" |  |  | *Anopheles gambiae* "VK" |  |  |
| --- | --- | --- | --- | --- | --- | --- | --- | --- |
|  |  |  | KDT<sub>50</sub> (min) | KDT<sub>95</sub> (min) | Mortality (%) | KDT<sub>50</sub> (min) | KDT<sub>95</sub> (min) | Mortality (%) |
| C1 | C.n 10% : O.a 90% | 0.1 | 102 | 132.9 | 3.61 | 171.6 | 321.3 | 2.53 |
|  |  | 0.5 | 9.6 | 22.1 | 18.49 | 33 | 57.3 | 16.17 |
|  |  | 1 | 4.4 | 13.2 | 29.79 | 27.4 | 50.3 | 19.78 |
| C2 | C.n 20% : O.a 80% | 0.1 | 131.7 | 232.7 | 1.94 | 222 | 310 | 1.15 |
|  |  | 0.5 | 3.3 | 13.2 | 32.29 | 35 | 73.3 | 13.25 |
|  |  | 1 | 2.4 | 10.2 | 68.97 | 16 | 25.9 | 26.08 |
| C3 | C.n 30% : O.a 70% | 0.1 | 142.8 | 232.7 | 6.66 | 178.1 | 304 | 6.25 |
|  |  | 0.5 | 10.8 | 21 | 32.04 | 25 | 38.5 | 13.68 |
|  |  | 1 | 5.5 | 13.4 | 91.67 | 16 | 16.1 | 20.21 |
| C4 | C.n 40 % : O.a 60 % | 0.1 | 108.7 | 147.1 | 22.06 | 142.4 | 172.1 | 4.1 |
|  |  | 0.5 | 7.3 | 16.9 | 78.49 | 20.3 | 47.6 | 17.11 |
|  |  | 1 | 6.7 | 12.4 | 98.78 | 7.7 | 14.3 | 42.39 |
| C5 | C.n 50% : O.a 50% | 0.1 | 91 | 153.6 | 4.55 | 106.7 | 182.8 | 3.66 |
|  |  | 0.5 | 11.2 | 18.6 | 28.89 | 23.5 | 78.7 | 19.48 |
|  |  | 1 | 0.5 | 13.1 | 97.78 | 7.1 | 16.5 | 30.38 |
| C6 | C.n 60% : O.a 40% | 0.1 | 77.4 | 145 | 19.39 | 116.1 | 168.1 | 7.08 |
|  |  | 0.5 | 15.7 | 37.6 | 91.67 | 48.1 | 64.9 | 22.38 |
|  |  | 1 | 8.6 | 13.4 | 98.9 | 8.9 | 20.7 | 45.21 |
| C7 | C.n 70% : O.a 30% | 0.1 | 131.9 | 245.3 | 3.49 | 166.1 | 303.3 | 1.19 |
|  |  | 0.5 | 48.7 | 99.6 | 17.84 | 137.2 | 225 | 15.21 |
|  |  | 1 | 11.6 | 17.4 | 37.98 | 13.2 | 28.8 | 19.62 |
| C8 | C.n 80% : O.a 20% | 0.1 | 76.4 | 150.6 | 15.12 | 119 | 218.2 | 8.75 |
|  |  | 0.5 | -25.2 | 28.8 | 95.65 | 24.2 | 52.7 | 27.38 |
|  |  | 1 | -26.9 | 1.4 | 100 | -2.1 | 12.2 | 80.7 |
| C9 | C.n 90% : O.a 10% | 0.1 | 23.5 | 77 | 13.98 | 53.5 | 126.7 | 12.15 |
|  |  | 0.5 | -32.3 | 3.7 | 96.7 | -5 | 7.4 | 94 |
|  |  | 1 | -227.9 | -70.1 | 100 | -227.3 | -69.6 | 100 |
|  | Permethrin | 0,75 | 16.3 | 30 | 100 | 41.2 | 90.4 | 62.5 |

The lowest rates of mortality were obtained with C7 (C.n 70% : O.a 30%) giving values of 37.98 and 19.62% at the higher concentration (1%) on Kisumu and VK strains, respectively (Table 2). On the VK strain, the LC50 values of the combinations C9, C8, C6 and C4 were less than 0.5% except for C5, C3 and C2. On the two *An. gambiae* strains, LC 50 values were significantly different for all combinations.

### Effects of combinations against *Anopheles gambiae*

The interactive effects between the two EOs in the binary combinations (Table 4) were obtained by the fractional inhibitory concentration (FIC) indices.

**Table 4:** LC_50_, LC_99_ and diagnostic concentration for all essential oils and combinations of *C. nardus* and *O. americanum* tested on *Anopheles gambiae* populations

| HE/<br>Combinations | Kisumu |  | VK |  | Diagnostic<br>concentration<br>(%) | Diagnostic<br>concentration<br>(mg/ml) | Diagnostic<br>concentration<br>mg/cm<sup>2</sup> |
| --- | --- | --- | --- | --- | --- | --- | --- |
|  | CL50(%)<br>(CL95) | CL99(%)<br>(CL95) | CL50(%)<br>(CL95) | CL99(%)<br>(CL95) |  |  |  |
| *C. citratus* | 2.3<br>(1.9-2.7) | 4.7<br>(4.4-6) | 2.7<br>- | 4.84<br>- | 9.4 | 94 | 1.04 |
| *C. nardus* | 0.63<br>(0.6-0.7) | 1.66<br>(1.4-2) | 1.12<br>(1-1.3) | 2.16<br>(1.8-2.7) | 3.32 | 33.2 | 0.37 |
| *E. camaldulensis* | 2.86<br>(1.6-9.3) | 5.26<br>(4.1-5.5) | 2.94<br>- | 5.05<br>- | 10.1 | 101 | 1.12 |
| *L. multiflora* | 0.21<br>0.16-0.3 | 0.74<br>(0.6-0.9) | 0.67<br>(0.6-0.7) | 1.17<br>(1.1-1.3) | 1.48 | 14.8 | 0.16 |
| *O. americanum* | 2.18<br>(1.5-3) | 4.13<br>(2.6-5.5) | 2.21<br>(1.5-3.5) | 4.59<br>(2.9-5.8) | 9.18 | 91.8 | 1.02 |
| C1 (C.n 10% : O.a 90%) | 1.37<br>(1.1-1.8) | 2.95<br>(2.3-3.4) | 2.9<br>- | 6.4<br>- | 5.9 | 59 | 0.66 |
| C2 (C.n 20% : O.a 80%) | 0.76<br>(0.7-0.8) | 1.69<br>(1.5-2) | 1.18<br>(1.0-1.4) | 2.45<br>(2-3.3) | 3.38 | 33.8 | 0.38 |
| C3 (C.n 30% : O.a 70%) | 0.61<br>(0.5-0.7) | 1.31<br>(1.2-1.5) | 2.07<br>(1.5-5) | 5<br>(4.2-5.9) | 2.62 | 26.2 | 0.29 |
| C4 (C.n 40% : O.a 60%) | 0.46<br>(0.4-0.5) | 1.19<br>(1-1.4) | 1.25<br>(1.2-1.5) | 2.48<br>(2-3.4) | 2.38 | 23.8 | 0.26 |
| C5 (C.n 50% : O.a 50%) | 0.57<br>(0.5-0.6) | 1.16<br>(1-1.3) | 1.36<br>(1.1-2) | 3.26<br>(2.4-5.5) | 2.32 | 23.2 | 0.26 |
| C6 (C.n 60% : O.a 40%) | 0.28<br>(0.2-0.3) | 1<br>(0.9-1.2) | 1.08<br>- | 2.15<br>- | 2 | 20 | 0.22 |
| C7 (C.n 70% : O.a 30%) | 1.93<br>(1.4-4) | 4.3<br>(3-4.7) | 2.44<br>(1.6-3.2) | 5.07<br>(3.1-5.6) | 8.6 | 86 | 0.96 |
| C8 (C.n 80% : O.a 20%) | 0.23<br>(0.2-0.3) | 0.58<br>(0.5-0.7) | 0.89<br>(0.8-1) | 1.3<br>(1.9-3) | 1.16 | 11.6 | 0.13 |
| C9 (C.n 90% : O.a 10%) | 0.22<br>(0.2-0.3) | 0.52<br>(0.4-0.6) | 0.32<br>(0.3-0.4) | 0.59<br>(0.5-0.7) | 1.04 | 10.4 | 0.12 |

LC = Lethal concentration ; CL = confidence limits ; Cn = *Cymbopogon nardus*; Oa = *Ocimum americanum*

**Table 5:** Effects of combinations of essential oils *Cymbopogon nardus* (C.n) and *Ocimum americanum* (O.a) on adults of *Anopheles gambiae* strain Kisumu and type of interactions (n = 125 adult).

| Combination of<br>essential oil (%) | Codes | LC<sub>50</sub><br>% | FIC<sub>Cn</sub> | FIC<sub>O.a</sub> | FIC | Effect |
| --- | --- | --- | --- | --- | --- | --- |
| C.n 0% : O.a 100% | - | 2.18 | - | - | - | - |
| C.n 10% : O.a 90% | C1 | 1.37 | 2.17 | 0.63 | 2.80 | No effect |
| C.n 20% : O.a 80% | C2 | 0.76 | 1.21 | 0.35 | 1.55 | No effect |
| C.n 30% : O.a 70% | C3 | 0.61 | 0.97 | 0.28 | 1.25 | No effect |
| C.n 40% : O.a 60% | C4 | 0.46 | 0.73 | 0.21 | 0.94 | Addition |
| C.n 50 % : O.a 50% | C5 | 0.57 | 0.90 | 0.26 | 1.17 | No effect |
| C.n 60 % : O.a 40% | C6 | 0.28 | 0.44 | 0.13 | 0.57 | Additive |
| C.n 70% : O.a 30% | C7 | 1.93 | 3.06 | 0.89 | 3.95 | No effect |
| C.n 80% : O.a 20% | C8 | 0.23 | 0.37 | 0.11 | 0.47 | Synergistic |
| C.n 90% : O.a 10% | C9 | 0.22 | 0.35 | 0.10 | 0.45 | Synergistic |
| C.n 100% : O.a 0% | - | 0.63 | - | - | - | - |

FIC = FIC<sub>Cn</sub> + FIC<sub>O.a</sub>; Synergistic: FIC < 0.5; Additive; 0.5 ≤ FIC ≤ 1; No effect: 1 ≤ FIC ≤ 4; Antagonistic: FIC > 4.

**Table 6:** Effects of combinations of essential oils *Cymbopogon nardus* (C.n) and *Ocimum americanum* (O.a) on adults of *Anopheles gambiae* VK strain and type of interaction (n = 125 adult).

| Combination of essential oil (%) | Codes | LC<sub>50</sub><br>% | FIC<sub>Oa</sub> | FIC<sub>Nc</sub> | FIC | Effect |
| --- | --- | --- | --- | --- | --- | --- |
| C.n 0% : O.a 100% | Oa | 2.21 | - | - | - | - |
| C.n 10% : O.a 90% | C1 | 2.9 | 1.31 | 2.59 | 3.9 | No effect |
| C.n 20% : O.a 80% | C2 | 1.18 | 0.53 | 1.05 | 1.58 | No effect |
| C.n 30% : O.a 70% | C3 | 2.07 | 0.94 | 1.85 | 2.79 | No effect |
| C.n 40% : O.a 60% | C4 | 1.25 | 0.56 | 1.12 | 1.68 | No effect |
| C.n 50 % : O.a 50% | C5 | 1.36 | 0.61 | 1.2 | 1.81 | No effect |
| C.n 60 % : O.a 40% | C6 | 1.08 | 0.49 | 0.96 | 1.45 | No effect |
| C.n 70% : O.a 30% | C7 | 2.44 | 1.1 | 2.17 | 3.27 | No effect |
| C.n 80% : O.a 20% | C8 | 0.89 | 0.40 | 0.79 | 1.19 | No effect |
| C.n 90% : O.a 10% | C9 | 0.32 | 0.14 | 0.28 | 0.42 | Synergistic |
| C.n 100% : O.a 0% | Cn | 1.12 | - | - | - | - |

FIC = FIC<sub>Cn</sub> + FIC<sub>Oa</sub>; Synergistic: FIC < 0.5; Addition; 0.5 ≤ FIC ≤ 1; No effect: 1 ≤ FIC ≤ 4; Antagonistic: FIC > 4.

These combination effects were classified as synergistic, additive, indifferent or antagonistic. In the present study, they varied according to the strain of *An. gambiae*. The combinations that showed synergistic and additive effects were also those were more effective in terms of mortality. Among the nine (9) binary combinations done and tested against *An. gambiae*, only C9 and C8 showed a synergistic effect while C6 and C4 showed an additive effect in Kisumu populations (Table 4). However, the effect of C6 was close to the synergistic effect (FIC_C6_ = 0.57). The other five combinations had no effect. On VK population, only C9 showed a synergistic effect.

### Diagnostic concentrations

Table 3 summarizes the lethal concentration for 50 and 99% mortality (LC_50_ and LC_99_) expressed with 95% confidence limits and diagnostic concentrations for all essential oils and all combinations prepared with EOs of *C. nardus* and *O. americanum*. The diagnostic concentration was obtained from twice the LC_99_ on susceptible strain ^3^.

The lowest diagnostic concentrations of 1.04%, 1.16% and 1.48% were obtained with C9 and C8, and *L. multiflora* OE, respectively. These three diagnostic concentrations were not significantly different from the diagnostic dose of permethrin (0.75%), the positive control. The highest diagnostic doses were obtained with C1 (5.9%), C7 (8.6%), *O. americanum* (9.18%), *C. citratus* (9.4%) *and camaldulensis* (10.1%).

## DISCUSSION

Up to now, the management of insecticide resistance remains a major challenge to achieving effective malaria elimination^1^. Indeed, pyrethroid resistance has been reported in 27 sub-Saharan African countries, raising the urgency of finding alternatives to these insecticides^25,26^. Searching for alternatives to chemical insecticides based on plant extracts may constitute new avenues for controlling malaria mosquito vectors populations.

Here, we evaluate the toxicity level of *Cymbopogon citratus, Cymbopogon nardus, Eucalyptus camaldulensis, Lippia multiflora* and *Ocimum americanum* EOs. Also, the binary combinations of *Cymbopogon nardus* (C.n) and *Ocimum americanum* (O.a) were examined in terms of toxicity.

Overall, almost all the EOs tested have shown an adulticidal effect on the susceptible (Kisumu) and on the resistant (VK) strain of *An. gambiae*. This insecticidal effect, which is highlighted through knock-down effects, lethal concentrations (LC) and rates of mortality (%) of the adult mosquitoes tested, varied significantly according to the concentrations and kind of EOs used. Among the EO tested, that of *L. multiflora* remains the most toxic on the two strains of *An. gambiae* tested followed by *C. nardus*. The least toxic EO was *E. camaldulensis*. Interestingly, *L. multiflora* may constitute a good alternative regarding the rate of mortality reaching 96.88%. The toxicity of EO of *L. multiflora* in the current study was significantly better than that of *Lantana camara, Hyptis spicigera, Hyptis suaveolens* EOs evaluated on *An. gambiae* strains from Kisumu and fields by Wangrawa et al^27^.

The difference observed in toxicity between the EOs tested in our investigations could be explained by their chemical composition. Indeed, several previous studies had shown that the bioactivity of an EO is attributed to the major compounds as they may constitute the most important part of the total compounds of the EO^16,28,29^. Hence, the toxicity of *L. multiflora* could be explained by its major compounds which are ß caryophyllene, p-cymen, thymol acetate and 1.8 cineol. According to previous studies done by Bassolé et al.^30^, the toxicity of *L. multiflora* EO on *Anopheles gambiae* and *Aedes aegypti* larvae was due to the presence of three major components: thymol, p-cymene and thymol acetate. *Hyptis suaveolens* EO containing also β-caryophyllene and 1.8 cineol^31^ has a high insecticidal activity^32^. Thus, the high insecticidal activity of *L. multiflora* EO was attributed to β-caryophyllene and 1.8 cineol which are major compounds, in the current study. The low insecticidal activity of *E. camaldulensis* EO could be explained probably by the absence of β-caryophyllene. Here, *C. nardus* EO do show intermediate adulticidal effect on *An. gambiae* populations tested unlike that found by Zulfikar^33^. According to these authors, the highest bioactivity from *C. nardus* was due to the presence of geraniol, the same compound found also in *C. nardus* tested in our current study. In fact, no effect was detected with *C. nardus*. The EOs which were less efficient as adulticides in the current study could be efficient as repellents. Indeed, previous studies shown the repellent effect of these EO on mosquitoes^34,35,36^. The KDT values of these EO mainly at the 1% concentration could explain this repellent property highlighted by irritant activity.

In addition to their adulticide activity, *L. multiflora* exhibits also toxic effect on larval populations of *An. gambiae* from the same locality^37^.

Overall, *C. nardus* and *O. americanum* oils provide intermediate rate of mortality. Do their combinations may improve the bioefficiency against *An. gambiae* populations? For this purpose, combinations from C1 to C9 were made, each concentration combining a certain proportion of each EO.

Globally, all combinations of the 2 EOs have improved the overall efficiency of the EOs compared to individual EOs. In the current study, the improvement of the adulticidal potential by the combination of the two EOs depends on their ratio in the combination. Indeed, combinations containing 60%, 80% and 90% of *C. nardus* EO were more effective than *C. nardus* EO tested alone. This is in agreement with the work of Bekele and Hassanali^38^; Pavela^39^ who had reported that the biological activity of essential oils depends not only on their qualitative composition, but also on the quantitative ratio of their constituents.

Improved efficiency was observed when knock down times were reduced and also synergistic and additive effects were observed for the combinations where the proportions of *C. nardus* reach 90%, 80% and 60% and 40%, respectively. This improved toxicity by the combinations of the 2 EOs, could be explained by the combined toxic effect of the major compounds. Previous works have showed that the toxic action of essential oils is due to the combined effects of different components, with or without significant individual toxic action against insects^40,41^. According to Burt^42^, individual EOs contain complex components that, when combined with each other, can lead to indifferent, additive, synergistic or antagonistic effects. In an earlier study, Abbassy et al^43^ reported that, in some cases, the whole EO may have a higher insecticidal activity than its isolated major components. For these authors, the minor compounds were essential for the bioefficiency and could allow a synergistic effect or potential influence.

The combinations showing a synergistic or additive effect would have, on one hand, both some major compounds of the EOs of *C. nardus* and *O. americanum* coupled with a variety of minor compounds. Previous studies have shown that the bioactivity of the essential oil is a consequence of interaction between the major components, but also other compounds, eventually oligoelements explaining combined effects, additive action between chemical classes and synergism or antagonism^30,44,45^. Generally, it seems that the effect of an active compound can be boosted by other major compounds and/or stimulated by minor compounds to give additive or synergistic effects^46^. Therefore, according to Berenbaum and Neal^47^, minor components present in low percentages can act as synergists, enhancing the bioefficiency of major constituents by various mechanisms.

## CONCLUSION

The current study confirmed that the essential oils of *Cymbopogon citratus, Cymbopogon nardus, Eucalyptus camaldulensis, Lippia multiflora* and *Ocimum americanum* have insecticidal properties. *Lippia multiflora* was efficient in comparison with the others tested. Our current study showed that the activity of the two combined essential oils, *Cymbopogon nardus* and *Ocimum americanum*, was improved by the combinations at certain proportions regarding the values of rate of mortality reaching at least 80%.

The essential oil of *Lippia multiflora* and combinations of EO of *C. nardus* and *O. americanum* could be valuable alternatives in the malaria vector control.

## DATA AVAILABILITY STATEMENT

The datasets generated and analyzed during the current study are available from the corresponding author on reasonable request. Requests to access these datasets should be directed to the corresponding author.

## AUTHOR CONTRIBUTIONS

MB and OG designed the study. DDS critically supervised the study. MB, IS, HK, and GBM carried out the laboratory experiments. MB, FSD and OG analyzed and interpreted the data and drafted the manuscript. OG, RKD, HCRN, MN, IHNB, revised the manuscript. All authors contributed to the article and approved the submitted version.

## ACKNOWLEDGMENTS

We are indebted to “Institut de recherche en Sciences Appliquées et technologiques” (IRSAT) for providing us with essential oils. I also thank Casimir Gnankiné for editing this manuscript.

## REFERENCES

1. WHO. Global plan for insecticide management. 130 (2012).

2. WHO, 2020. WHO, 30 novembre 2020. vol. 2507 (2020).

3. OMS. Procédures pour tester la résistance aux insecticides chez les moustiques vecteurs du paludisme Seconde édition. (2017).

4. WHO. Guidelines for Malaria Vector Control. Guidelines for Malaria Vector Control (2019).

5. Churcher, T. S., Lissenden, N., Griffin, J. T., Worrall, E. & Ranson, H. The impact of pyrethroid resistance on the efficacy and effectiveness of bednets for malaria control in Africa. Elife 5, (2016).

6. Hemingway, J. et al. Averting a malaria disaster: will insecticide resistance derail malaria control? The Lancet 387, 1785–1788 (2016).

7. Dabiré1 et al. Trends in Insecticide Resistance in Natural Populations of Malaria Vectors in Burkina Faso, West Africa: 10 Years’ Surveys K. Intech 32,.479-502, (2012).

8. WHO. WHO Global Malaria Programme: Global Plan For Insecticide Resistance Management. (2012).

9. Toe, K. H. et al. Do bednets including piperonyl butoxide offer additional protection against populations of Anopheles gambiae s.l. that are highly resistant to pyrethroids? An experimental hut evaluation in Burkina Fasov. Med. Vet. Entomol. 32, 407–416 (2018).

10. BayiliI K. et al. Experimental hut evaluation of DawaPlus 3. 0 LN and DawaPlus 4. 0 LN treated with deltamethrin and PBO against free-flying populations of Anopheles gambiae s. l. in Valle du Kou, Burkina Faso. PLoS One 14, 1–15 (2019).

11. Hien, A. S. et al. Evidence supporting deployment of next generation insecticide treated nets in Burkina Faso: bioassays with either chlorfenapyr or piperonyl butoxide increase mortality of pyrethroid-resistant Anopheles gambiae. Malar. J. 20, 1–12 (2021).

12. Zoubiri, S. & Baaliouamer, A. Potentiality of plants as source of insecticide principles. J. Saudi Chem. Soc. 18, 925–938 (2014).

13. Tripathi, A. K., Upadhyay, S., Bhuiyan, M. & Bhattacharya, P. R. A review on prospects of essential oils as biopesticide in insect-pest management. J. of Pharmacognosy Phytother 1, 52–63 (2009).

14. Isman, M. B. Plant essential oils for pest and disease management. Crop Prot. 19, 603–608 (2000).

15. Mossa, A. T. H. Green Pesticides: Essential oils as biopesticides in insect-pest management. J. Environ. Sci. Technol. 9, 354–378 (2016).

16. Lucia, A. et al. Larvicidal effect of Eucalyptus grandis essential oil and turpentine and their major components on aedes Aegypti larvae. J. Am. Mosq. Control Assoc. 23, 299–303 (2007).

17. Singh, R., Koul, O. & Rup, P. J. Toxicity of some essential oil constituents and their binary mixtures against Chilo partellus (Lepidoptera□: Pyralidae). International J. of Tropical Insect Science 29, 93–101 (2009).

18. Sarma, R., Adhikari, K., Mahanta, S. & Khanikor, B. Combinations of Plant Essential Oil Based Terpene Compounds as Larvicidal and Adulticidal Agent against Aedes aegypti (Diptera: Culicidae). Sci. Rep. 9, 1–13 (2019).

19. Mansour, S. A., Foda, M. S. & Aly, A. R. Mosquitocidal activity of two Bacillus bacterial endotoxins combined with plant oils and conventional insecticides. Ind. Crops Prod. 35, 44–52 (2012).

20. Yaméogo, F., Wendgida, D., Sombié, A., Sanon, A. & Badolo, A. Insecticidal activity of essential oils from six aromatic plants against Aedes aegypti, dengue vector from two localities of Ouagadougou, Burkina Faso. Arthropod. Plant. Interact. 15, 627–634 (2021).

21. Raphael N’Guessan, Vincent Corbel, Martin Akogbéto, and M. R. treated Nets and Indoor Residual Reduced Efficacy of Insecticide-Pyrethroid Resistance Area, Benin. Emerg. Infect. Dis. 13, 199–206 (2007).

22. Abbott, W. S. A method of computing the effectiveness of an insecticidel. journal of the arunnrclu moseurro coxrnol AssocrATroN Vot vol. 3 (1925).

23. Schelz, Z., Molnar, J. & Hohmann, J. Antimicrobial and antiplasmid activities of essential oils. Fitoterapia 77, 279–285 (2006).

24. Bassolé, I. H. N. & Juliani, H. R. Essential oils in combination and their antimicrobial properties. Molecules 17, 3989–4006 (2012).

25. Tchoumbougnang, F. et al. Activité larvicide sur Anopheles gambiae Giles et composition chimique des huiles essentielles extraites de quatre plantes cultivées au Cameroun. Biotechnol. Agron. Soc. Environ. 13, 77–84 (2009).

26. Ranson, H. & Lissenden, N. Insecticide Resistance in African Anopheles Mosquitoes: A Worsening Situation that Needs Urgent Action to Maintain Malaria Control. Trends Parasitol. 32, 187–196 (2016).

27. Wangrawa, Di. W. et al. Insecticidal Activity of Local Plants Essential Oils Against Laboratory and Field Strains of Anopheles gambiae s. L. (Diptera: Culicidae) from Burkina Faso. J. Econ. Entomol. 111, 2844–2853 (2018).

28. Gbolade, A. A. & Lockwood, G. B. Toxicity of Ocimum sanctum L. Essential oil to Aedes aegypti Larvae and its Chemical Composition. J. Essent. Oil-Bearing Plants 11, 148–153 (2008).

29. Raseetha Vani, S., Cheng, S. F. & Chuah, C. H. Comparative study of volatile compounds from genus Ocimum. Am. J. Appl. Sci. 6, 523–528 (2009).

30. Bassolé IHN, Guelbéogo WM, Nébié R, Costantini C, Sagnon NF, Kabore ZI, T. S. Ovicidal and larvicidal activity against Aedes aegypti and Anopheles gambiae complex mosquitoes of essential oils extracted from three spontaneous plants of Burkina Faso. Parasitologia 45□:23–26. Parasitologia 45, 23–26 (2003).

31. MarcotuN. Peerzadallio. Chemical Composition of the Essential Oil of Hyptis Suaveolens. Molecules 2, 165−168 (1997).

32. Ilboudo, Z. et al. Biological activity and persistence of four essential oils towards the main pest of stored cowpeas, Callosobruchus maculatus (F.) (Coleoptera: Bruchidae). J. Stored Prod. Res. 46, 124–128 (2010).

33. Zulfikar Aditama, W. & Sitepu, F. Y. The effect of lemongrass (Cymbopogon nardus) extract as insecticide against Aedes aegypti. Int. J. Mosq. Res. 6, 101–103 (2019).

34. Ojewumi M.E., Adeyemi A.O. & Ojewumi E.O. Oil Extract From Local Leaves -an Alternative To Synthetic Mosquito Repellants. Pharmacophore 9, 1–6 (2018).

35. Gnankiné, O. & Bassolé, I. L. H. N. Essential oils as an alternative to pyrethroids’ resistance against anopheles species complex giles (Diptera: Culicidae). Molecules 22, (2017).

36. Bossou, A. D. et al. Chemical composition and insecticidal activity of plant essential oils from Benin against Anopheles gambiae (Giles). Parasit. Vectors 6, 337 (2013).

37. Balboné M. et al. Essential oils from five local plants: An alternative. Front. in Trop. Dis.3, 853405 (2022).

38. Bekele, J. & Hassanali, A. Blend effects in the toxicity of the essential oil constituents of Ocimum kilimandscharicum and Ocimum kenyense (Labiateae) on two post-harvest insect pests. Phytochemistry 57, 385–391 (2001).

39. Pavela, R. Acute and synergistic effects of some monoterpenoid essential oil compounds on the house fly (Musca domestica l.). J. Essent. Oil-Bearing Plants 11, 451–459 (2008).

40. Tanprasit, P. Biological control of dengue fever mosquitoes (aedes aegypti linn.) using leaf extracts of chan (Hyptis suaveolens (l.) poit.) and hedge flower (Lantana camara linn.). Suranaree University of Technology, Nakhon ratchasima, Thailand (2005).

41. Park, H. M. et al. Larvicidal activity of myrtaceae essential oils and their components against Aedes aegypti, acute toxicity on Daphnia magna, and aqueous residue. J. Med. Entomol. 48, 405–410 (2011).

42. Burt, S. Essential oils: Their antibacterial properties and potential applications in foods - A review. International Journal of Food Microbiology. 94 223–253 (2004).

43. Abbassy, M. A., Abdelgaleil, S. A. M. & Rabie, R. Y. A. Insecticidal and synergistic effects of Majorana hortensis essential oil and some of its major constituents. Entomol. Exp. Appl. 131, 225–232 (2009).

44. Chiasson, H., Bélanger, A., Bostanian, N., Vincent, C. & Poliquin, A. Acaricidal properties of Artemisia absinthium and Tanacetum vulgare (Asteraceae) essential oils obtained by three methods of extraction. J. Econ. Entomol. 94, 167–171 (2001).

45. Luz T.R.S.A., de Mesquita L.S.S., Amaral F.M.M. do & Coutinho D.F. Essential oils and their chemical constituents against Aedes aegypti L. (Diptera: Culicidae) larvae. Acta Tropica. 212 (2020).

46. Emilie, D., Mallent, M., Menut, C., Chandre, F. & Martin, T. Behavioral Response of Bemisia tabaci (Hemiptera: Aleyrodidae) to 20 Plant Extracts. J. Econ. Entomol. 108, 1890–1901 (2015).

47. Berenbaum M.A.Y. & Neal J.J. Synergism between myristicin and xanthotoxin, a naturally cooccurring plant toxicant. Journal of Chemical Ecology 11, 1349–1358 (1985).

